# Interhemispheric transfer of working memories

**DOI:** 10.1101/2020.08.12.248203

**Authors:** Scott L. Brincat, Jacob A. Donoghue, Meredith K. Mahnke, Simon Kornblith, Mikael Lundqvist, Earl K. Miller

## Abstract

Visual working memory (WM) storage is largely independent between the left and right visual hemifields/cerebral hemispheres, yet somehow WM feels seamless. We studied how WM is integrated across hemifields by recording neural activity bilaterally from lateral prefrontal cortex. An instructed saccade during the WM delay shifted the remembered location from one hemifield to the other. Before the shift, spike rates and oscillatory power showed clear signatures of memory laterality. After the shift, the lateralization inverted, consistent with transfer of the memory trace from one hemisphere to the other. Transferred traces initially used different neural ensembles from feedforward-induced ones but they converged at the end of the delay. Around the time of transfer, synchrony between the two prefrontal hemispheres peaked in theta and beta frequencies, with a directionality consistent with memory trace transfer. This illustrates how dynamics between the two cortical hemispheres can stitch together WM traces across visual hemifields.

## INTRODUCTION

Imagine driving on the freeway. A car passes you and holds your attention as you wait to see if it will cut in front of you. Even if you briefly close your eyes or shift your gaze elsewhere, you are able to maintain the car’s location in mind and are surprised if it changes unexpectedly. This relies on visual working memory (WM), the ability to maintain images in mind in their absence. Decades of evidence points toward prefrontal cortex (PFC) as a key node in the cortical network underlying WM (D’Esposito and Postle, 2015; Funahashi et al., 1989; Fuster and Alexander, 1971; Miller et al., 1996; Romo et al., 1999; Ungerleider et al., 1998; Voytek and Knight, 2010).

In both humans and monkeys, visual WM seems largely independent between the left and right visual hemifields, which project to the right and left cerebral hemispheres, respectively. WM has a very limited storage capacity (Luck and Vogel, 1997, 2013). But the capacity within one visual hemifield is largely unaffected by the number of objects in the other hemifield (Buschman et al., 2011; Delvenne, 2005; Umemoto et al., 2010). Correspondingly, the neural correlates of WM storage and WM load (how many items are held in memory) primarily reflect items within the contralateral hemifield (Funahashi et al., 1990; Kastner et al., 2007; Kornblith et al., 2015; Luria et al., 2016; Rainer et al., 1998).

Nevertheless, visual cognition seems seamless across the visual field, even when eye movements switch the remembered location of objects between visual hemifields. In such situations, are memory representations transferred from one cerebral hemisphere to the other or are they “bound” to the hemisphere where they were initially stored? If transferred, do transferred WMs utilize the same neural representation as ones induced by feedforward visual inputs, and what are the mechanisms underlying transfer?

We utilized a novel variant of the delayed nonmatch-to-sample task. A midline-crossing saccade during the memory delay switched the hemifield of a remembered item. We found that WM traces are transferred from one prefrontal hemisphere to the other, transferred traces recruit neural ensembles distinct from those induced by feedforward inputs, and that this transfer is facilitated by rhythmic coupling between the cerebral hemispheres.

## RESULTS

Monkeys were trained to perform a modified version of a nonmatch-to-sample visual WM task (Fig. 1A). They fixated on a point on the left or right (50% of trials randomly) of a computer screen. An object briefly appeared as a sample in the center of the screen, thus in the right or left visual hemifield, respectively (Fig. 1B, left). The sample could be one of two different objects, at one of two different locations slightly above or below the center of the screen. The monkeys were required to remember both the object identity and location over a blank delay, and then compare it to a test object. If it did not match the sample in either identity or upper/lower location, they were trained to saccade directly to it. Otherwise, they held fixation until a second, always non-matching test object appeared.

**Figure 1.**
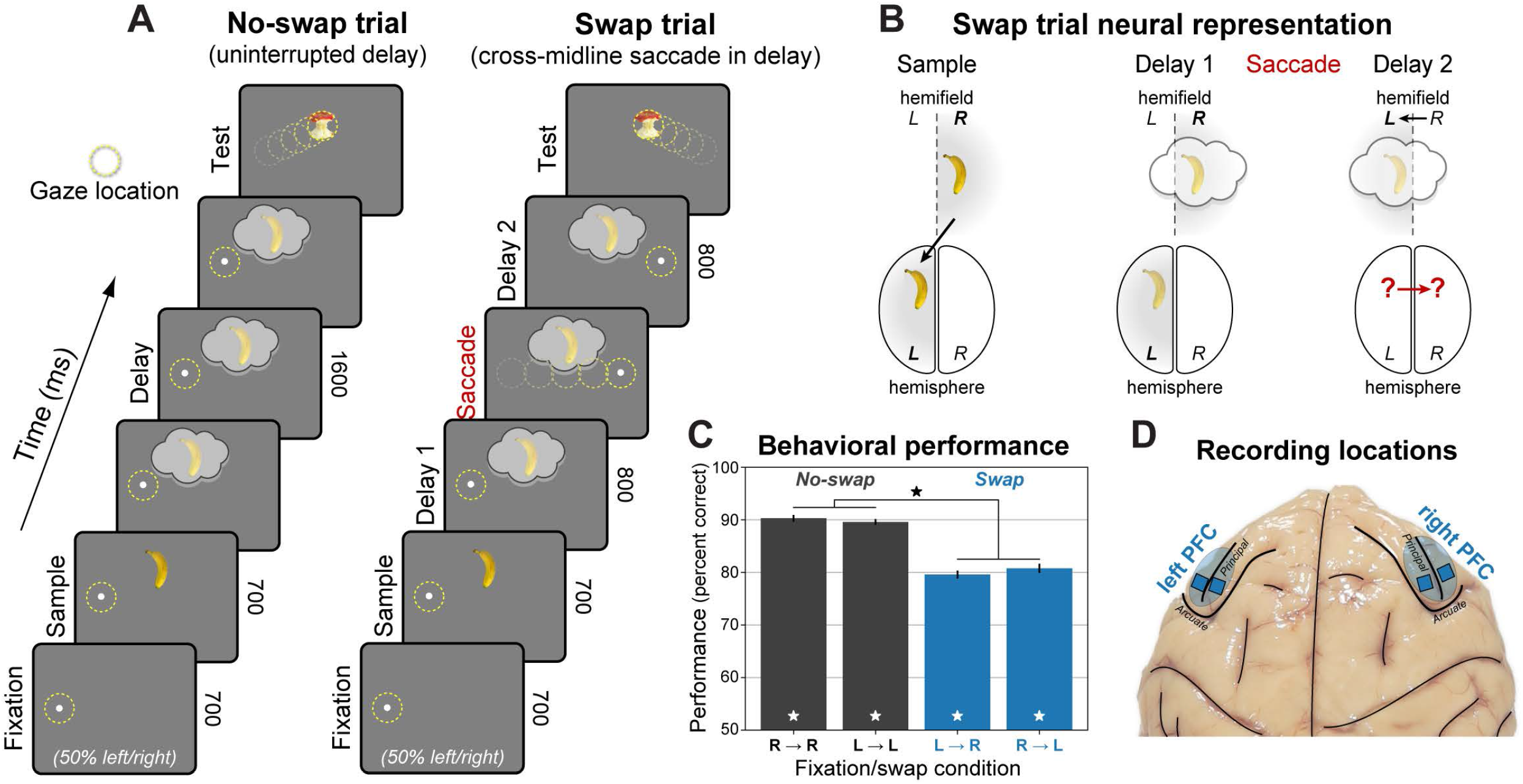
Behavioral and electrophysiological methods. (A) Hemifield-swap working memory (WM) task. Subjects fixated to the left or right while a sample object was presented in the center, placing it in the right or left visual hemifield, respectively. Samples could be one of two objects, presented in one of two locations (above or below center). Both the object and its location needed to be remembered over a blank 1.6 s delay. After the delay, a series of two test objects was displayed, and subjects responded to the one that did not match the sample in object identity or upper/lower location (response to first object shown for brevity). In “No-swap” trials (left), the WM delay was uninterrupted. In “Swap” trials (right), subjects were instructed to saccade to the opposite side mid-delay, switching the visual hemifield of the remembered location relative to gaze. (B) Neural representation of WM in Swap trials. The WM trace is initially encoded in the prefrontal hemisphere contralateral to the sample. We tested whether the change in gaze on Swap trials caused the WM traces to transfer to the other hemisphere. (C) Mean performance (± SEM across 56 sessions) for each swap condition and visual hemifield. Monkeys performed the task well (white stars: significant vs. chance), with a small but significant decrease (black star) on Swap trials. (D) Electrophysiological signals were recorded bilaterally from 256 electrodes in lateral prefrontal cortex (PFC).

A random 50% of trials had an uninterrupted delay, with no change in the remembered sample location relative to gaze (“*No-swap*” trials; Fig. 1A, left). On the other trials, halfway through the delay the fixation point jumped across the midline to the opposite side, instructing an immediate saccade and refixation on it for the remainder of the delay. This shifted the remembered sample’s retinotopic location to the opposite visual hemifield (“*Swap*” trials; Fig. 1A, right). Performance was good for all conditions (all *p* ≤ 1×10^−4^, randomized sign test across 56 sessions), albeit somewhat worse on Swap trials (*p* ≤ 1×10^−4^, permutation paired *t*-test; Fig. 1C). There were no significant differences in performance when the sample (*p* = 0.71) or test object (*p* = 0.10) appeared in the left vs. right hemifield, so all results were pooled across them. We recorded multi-unit activity (MUA) and local field potentials (LFPs) from 256 electrodes in four chronic arrays implanted bilaterally in both hemispheres of lateral prefrontal cortex during task performance (Fig. 1D). All presented statistics were based on nonparametric randomization tests across sessions, corrected for multiple comparisons across frequencies and time points (Benjamini and Yekutieli, 2001; see STAR Methods for details).

### Laterality of working-memory-related activity

We first examined No-swap trials to establish a reliable signature of the laterality of the working memory trace. Data from each prefrontal hemisphere and sample object hemifield was analyzed separately and then results were pooled based on whether the sample was contralateral or ipsilateral to the recorded hemisphere.

Spiking was stronger and more informative for contralateral than ipsilateral objects. Average MUA was significantly higher for contralateral than ipsilateral samples throughout the trial (Fig. 2A, stars; *p* < 0.01, corrected, permutation paired *t*-test). In fact, for ipsilateral samples, MUA was above baseline on average only during the response to the sample presentation and a brief “ramp-up” at the end of the delay (Fig. 2A, dots; *p* < 0.01, corrected, randomized sign test). Ipsilateral samples only had a weak effect on spiking during the delay. Unlike contralateral samples, which increased spiking, ipsilateral samples elicited a balance between increased and decreased spiking relative to baseline (Supplement Fig. S1).

**Figure 2.**
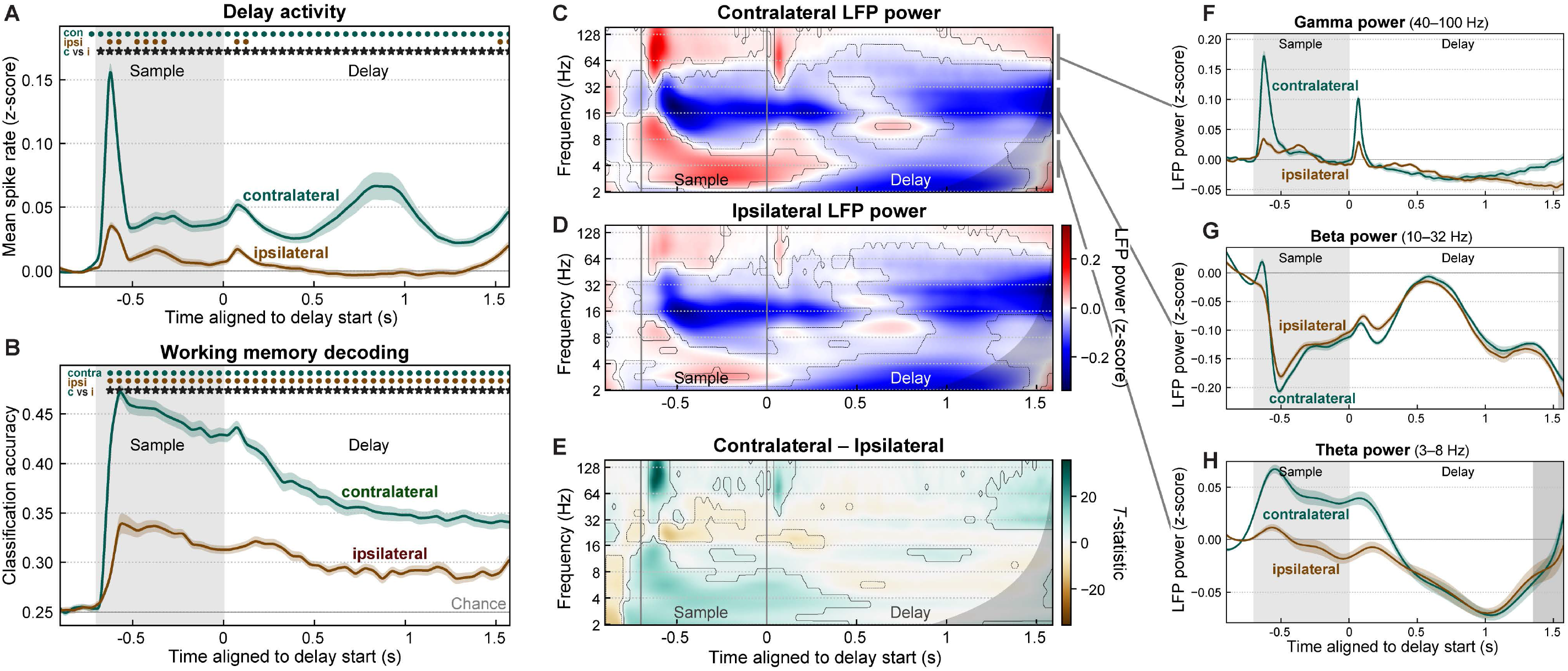
Contralateral bias in prefrontal cortex. (A) Population mean spike rates (multi-unit activity z-scored to baseline, ± SEM across 56 sessions) for sample objects contralateral (green) and ipsilateral (brown) to the recorded prefrontal hemisphere (pooled across left and right). Activity for contralateral samples was greater than baseline (green dots) and greater than activity for ipsilateral samples (stars). See also Figure S1. (B) Mean (± SEM) accuracy for decoding the item held in WM (object identity and upper/lower location) from prefrontal population spike rates, for samples in the contralateral (green) and ipsilateral (brown) visual hemifield. Contralateral decoding accuracy was greater than ipsilateral (stars). (C–D) Mean time-frequency LFP power (z-scored to baseline) for contralateral (C) and ipsilateral (D) samples. Contours indicate significant change from baseline. Gray regions to right of C–H indicate time points with possible temporal smearing of test-period effects. Gamma (∼40–100 Hz) and theta (∼3–8 Hz) power increased from baseline (red), while beta (∼10–32 Hz) power was suppressed from baseline (blue). (E) Contrast (paired-observation t-statistic map) between contralateral and ipsilateral power. Contours indicate significant difference. (F–H) Summary of LFP power for contralateral (green) and ipsilateral (brown) sample objects, pooled within frequency bands: gamma (F), beta (G), and theta (H). Due to off-axis structure in the time-frequency data, these 1D plots cannot fully capture it, and are thus included only to aid visualization. All modulations from baseline were stronger for contralateral samples, but only gamma showed a difference during the delay period. See also Figure S2.

We computed the accuracy of decoding the contents of WM as a measure of information conveyed in the neural population about WMs. We used a linear discriminant classifier to decode object identity and upper/lower location from the pattern of spiking activity across all MUAs within each hemisphere at each time point. Cross-validated decoding accuracy was significantly above chance for both contralateral and ipsilateral sample objects (Fig. 2B, dots; *p* < 0.01, sign test), but it was significantly higher for contralateral (Fig. 2B, stars; *p* < 0.01, paired *t*-test). Thus, prefrontal sensory and memory-related spiking showed a clear contralateral bias in strength and information content, as previously reported (see Discussion).

LFP power also exhibited a contralateral bias, especially for higher frequencies. Gamma power (∼40–100 Hz) was significantly elevated relative to baseline during sample object presentation and a “ramp-up” at the end of the delay for both contralateral (Fig. 2C) and ipsilateral (Fig. 2D) samples (*p* < 0.01, sign test; summarized in Fig. 2F). Gamma power induced by contralateral sample objects was significantly higher than for ipsilateral objects during the sample onset and offset transients and the pre-test “ramp-up” (Fig. 2E; *p* < 0.01, paired *t*-test). Theta power (∼3–8 Hz) also showed a contralateral bias during the sample response but not the memory delay (Fig. 2E,H). In contrast, beta power (∼10–32 Hz) showed effects in the opposite direction overall—significant *decreases* in power from baseline during the sample and late delay periods for both contralateral and ipsilateral sample objects (Fig. 2G). Like gamma and theta enhancement, beta suppression was significantly stronger for contralateral than ipsilateral samples (Fig. 2E), but, like theta power, only during the sample object response. Beta oscillatory burst rates exhibited a stronger and more sustained laterality than beta power, but analysis of LFP bursting produced otherwise similar results (Fig. S2A–B). These results indicate that prefrontal LFP power, like spiking activity, exhibits a clear contralateral bias.

### Transfer of working memories between cerebral hemispheres

We then leveraged the neural signature of WM laterality to examine what happened when the remembered location was shifted by a saccade to the opposite visual hemifield relative to the center of gaze. We propose two alternative hypotheses. On the one hand, a WM trace might be bound to the initial representation induced by visual inputs and might simply remain in the hemisphere where it was originally encoded (Fig. 3A). This *stable trace model* predicts no change in neural signatures after the mid-delay saccade (Fig. 3B). Alternatively, when the hemifield switches, the neural trace itself might also move from the hemisphere it was originally encoded in to the opposite hemisphere, now contralateral to the remembered location (Fig. 3C). This *shifting trace model* predicts neural signatures of laterality in the Swap trials will invert after the midline crossing saccade (Fig. 3D).

**Figure 3.**
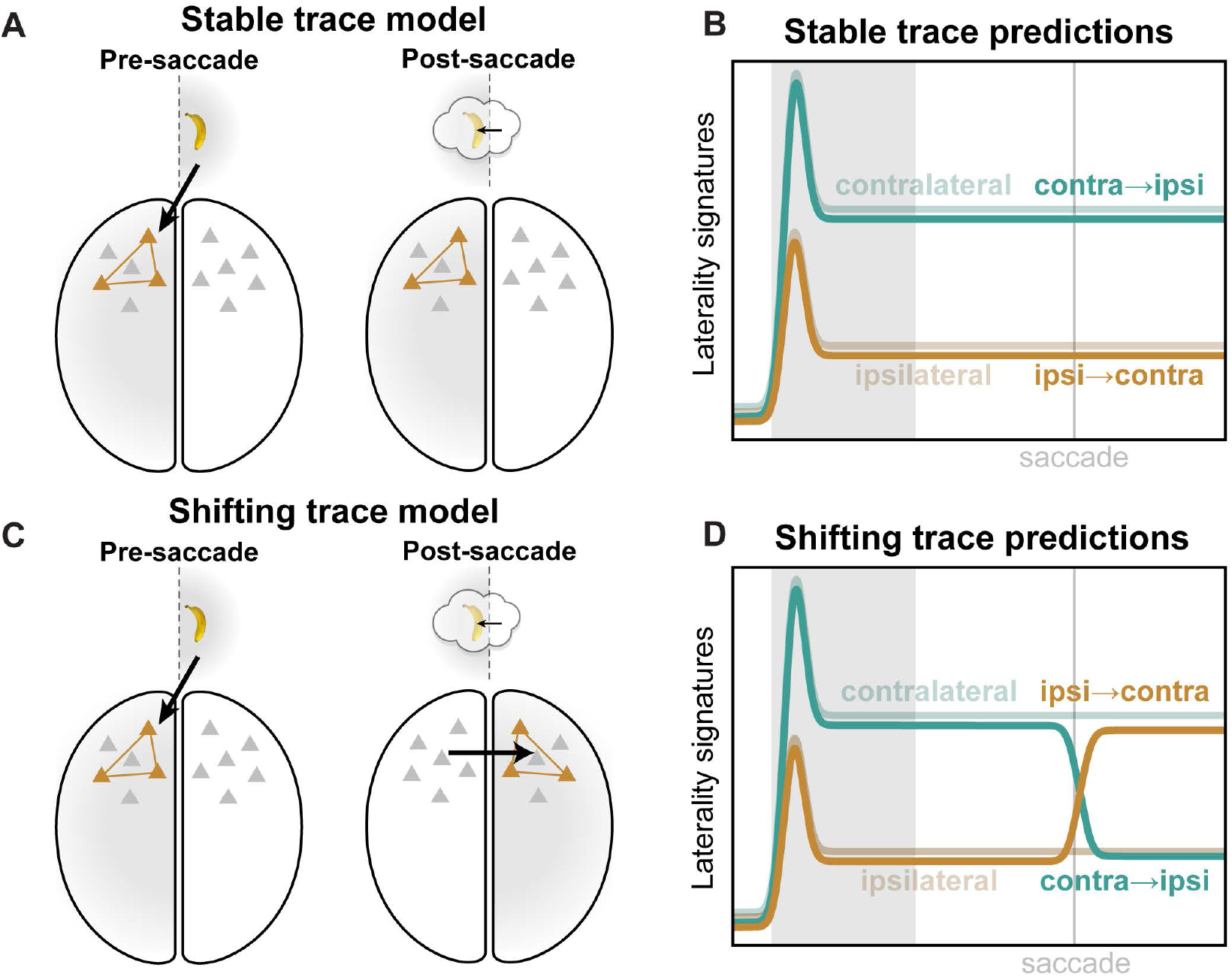
Competing models of swap effects. (A) The stable trace model posits that once a working memory is encoded in a given cortical hemisphere (left), it will remain there (right), despite the remembered location shifting from one hemifield to the other (inset). (B) This model predicts that neural signatures of memory trace laterality will be unaltered by the mid-delay saccade in our task. (C) The shifting trace model assumes that when the hemifield of the remembered location is swapped, the memory trace will be transferred from one cortical hemisphere to the other. (D) This model predicts a post-saccadic inversion of the neural signatures of laterality: shifting the remembered location into the contralateral hemifield (orange) should come to approximate the constant contralateral location (desaturated green), while shifting it ipsilateral (green) should come to look like constant ipsilateral trials (desaturated brown).

We found evidence for the shifting trace model, an inversion of neural laterality signatures after the midline-crossing saccade. This was apparent in average MUA rate (Fig. 4A) and in information carried by MUA (Fig. 4B). As in the No-swap trials (Fig. 2A,B), prefrontal MUA starts the delay with a bias toward the contralateral hemifield (Fig. 4A,B, left; ‘H’ symbols: *p* < 0.01, corrected, sample hemifield main effect in hemifield × shift condition permutation 2-way ANOVA). But after the saccade (Fig. 4A,B, right), both MUA and decoding accuracy increased for remembered locations that shifted from the ipsilateral to the contralateral hemifield (orange) relative to that for ipsilateral locations on No-swap trials (desaturated brown; brown stars: *p* < 0.01, paired *t*-test). By contrast, on Swap trials when the saccade shifted the remembered location from the contralateral to ipsilateral hemifield (green), there was a decrease compared to the contralateral location on No-swap trials (desaturated green; green stars: *p* < 0.01). For average MUA (Fig. 4A), there was a complete inversion. The spike rates after a saccade that shifted the remembered location into a given hemifield almost exactly matched those for a static memory in the same hemifield. Around the time of the saccade, there was also increased spiking for both Swap conditions, relative to the No-swap trials (‘S’ symbols: *p* < 0.01, shift condition main effect). Nevertheless, later in the delay, the predicted inversion effect was dominant (‘X’ symbols: *p* < 0.01, interaction effect). For decoding accuracy (Fig. 4B), ipsilateral-shifting trials (green) were near the value of constant ipsilateral trials (desaturated brown). Contralateral-shifting trials (orange) exhibited a bump of increased accuracy after the saccade (brown stars: *p* < 0.01, paired *t*-test) but subsequently declined and never attained the level of constant contralateral trials (desaturated green). This imperfect transfer of information may explain why behavioral performance was significantly decreased in Swap trials.

**Figure 4.**
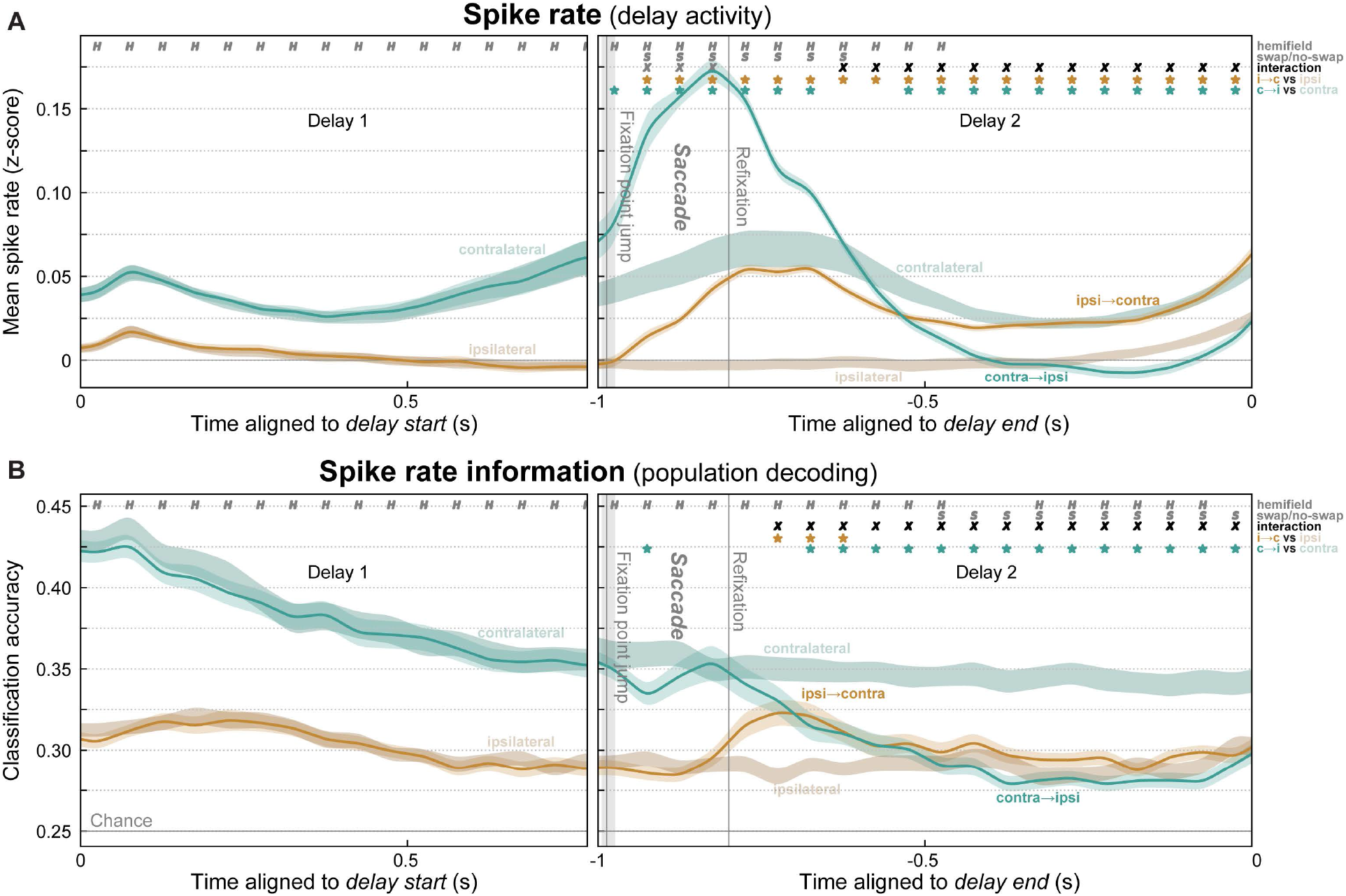
Evidence for interhemispheric transfer of working memory traces in prefrontal spiking activity. (A) Mean (± SEM) multi-unit spike rates for all trials where the remembered location was constant in the contralateral (desaturated green) or ipsilateral (desaturated brown) hemifield, or where it swapped from ipsilateral to contralateral (orange) or from contralateral to ipsilateral (green). Before the mid-delay saccade, there was only a significant effect of the sample hemifield (‘H’ symbols). Around the saccade, activity was greater overall for Swap than No-swap trials (‘S’ symbols). Later, the Swap trials inverted and approximated activity in the corresponding No-swap trials (‘X’ symbols: significant hemifield × swap condition interaction). Stars indicate significant difference of swap conditions from their respective No-swap baseline. (B) Mean (± SEM) accuracy for decoding the item held in WM from spike rates. As predicted, post-saccade accuracy decreased in contralateral-to-ipsilateral trials. Decoding on ipsilateral-to-contralateral trials also significantly improved above baseline, but only transiently, and it never reached the level of constant contralateral trials.

Similar effects were seen in LFP power (Fig. 5). During and just after the saccade, gamma (Fig. 5A–B, D) and theta (Fig. 5A–B,F) power were stronger overall for Swap than No-swap trials. Later in the delay, however, gamma power inverted and became stronger for remembered locations moving from the ipsilateral to the preferred contralateral hemifield (Fig. 5C,D; *p* < 0.01, hemifield × swap interaction effect) as predicted by the shifting trace model. Theta power showed a similar inversion at the very end of the delay, though this is likely due to temporal smearing of test period effects (Fig. 5C,F; gray regions indicate extent of possible test effects).

**Figure 5.**
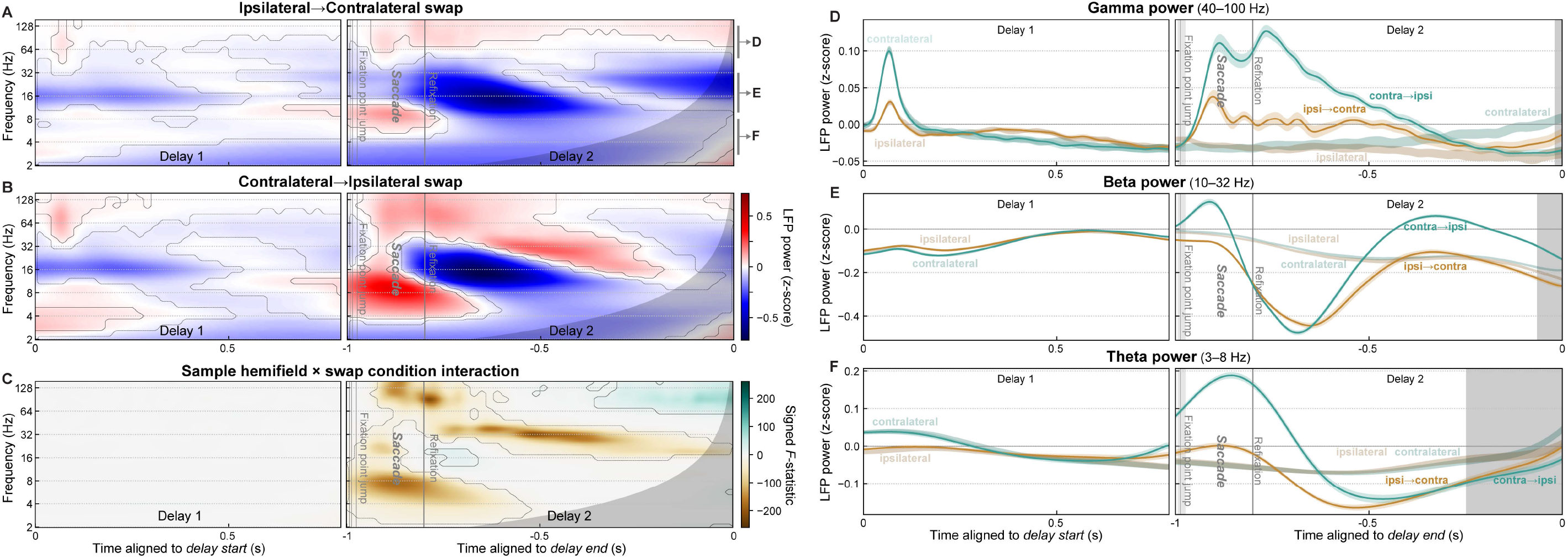
Evidence for interhemispheric transfer of working memory traces in prefrontal LFP power. (A–B) Mean time-frequency LFP power for trials where the remembered location shifted from the ipsilateral to contralateral (A) or from the contralateral to ipsilateral (B) hemifield. Contours indicate significant difference from pre-sample baseline. Gray regions at right of all panels indicate time points with possible temporal smearing of test-period effects. (C) Swap inversion effect. F-statistic map for sample hemifield × swap condition interaction, signed to indicate if power was greater when remembered location ends up in contralateral (green) or ipsilateral (brown) hemifield. Contours indicate significant interaction. (D–F) Summary of LFP power for ipsilateral-to-contralateral (green) and contralateral-to-ipsilateral (orange) trials, pooled within frequency bands labeled in panel A: gamma (D), beta (E), and theta (F). Around the time of the saccade, LFP power in all bands showed strong effects of the swap condition (saccade vs. no saccade). Later in the post-saccade delay, signatures in all bands inverted, as predicted by the shifting trace model. See also Figure S2.

Beta power (Fig. 5A–C,E) exhibited complex multiphasic dynamics on Swap trials (Fig. 5F). It was suppressed initially after the saccade, but later became enhanced, relative to power on No-swap trials (Fig. 5F). On top of these overall dynamics, however, beta power on the Swap conditions inverted—it became stronger for remembered locations shifting into the ipsilateral hemifield than for those shifting into the contralateral hemifield (Fig. 5C,F). Thus, as for spiking, prefrontal LFP power signatures of working memory laterality also exhibited the inversion predicted by the shifting trace model (Fig. 3D). These results support the hypothesis that the memory trace is transferred from one cortical hemisphere to the other.

### Interhemispheric transfer activates novel neural ensembles

A model consistent with the data thus far is that WMs transferred between hemispheres recruit the same neural ensembles as memory traces activated by feedforward visual inputs into the same cortical hemisphere. Under this *generic ensemble model* (Fig. 6A), when a given object— say, a banana in the upper location—is held in WM within a cortical hemisphere, it uses the same neural ensemble (i.e., the same pattern of spiking across our electrode arrays) whether it arrived there via feedforward inputs from visual cortex (Fig. 6A, left) or via interhemispheric inputs from the contralateral hemisphere (Fig. 6A, right). We could test this because the saccade on Swap trials brought the remembered sample location to the same retinotopic coordinates where it appeared on the No-swap trials. An alternative model is motivated by the fact that unique prefrontal ensembles are activated by different combinations of input features and task contexts (Rigotti et al., 2013). Perhaps the same information arriving via different circuits—feedforward (Fig. 6C, left) vs. interhemispheric (Fig. 6C, right)—also activates different ensembles. We call this alternative the *novel ensemble* model.

**Figure 6.**
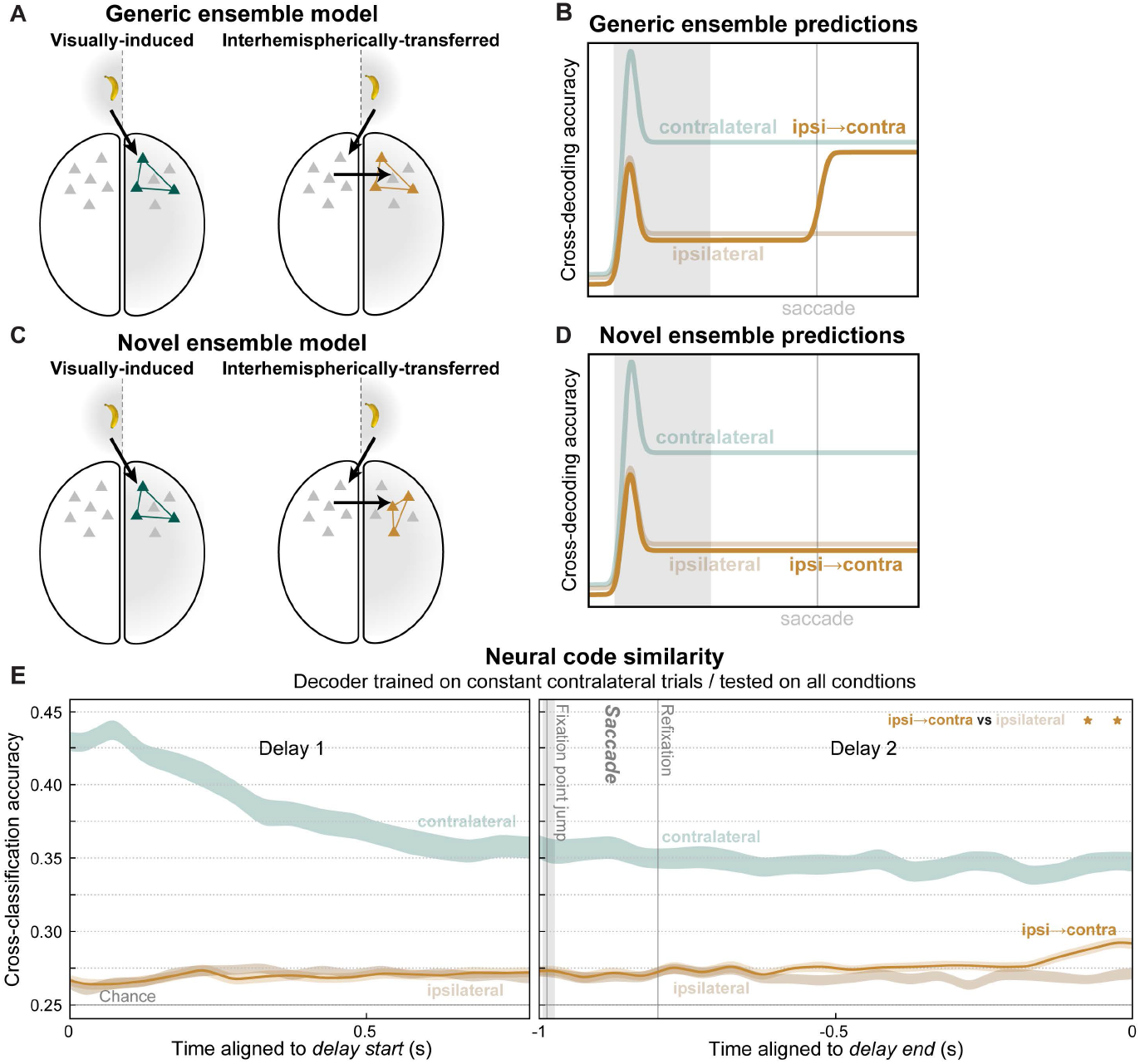
Transferred working memory traces utilized novel ensembles, but converged toward visually-induced ensembles at delay end. (A) The generic ensemble model assumes a given memory trace will activate the same neural ensemble (colored neurons) whether it arrives in prefrontal cortex via feedforward inputs from visual cortex (left) or via interhemispheric inputs from the opposite cortical hemisphere (right). (B) It predicts a classifier trained to decode working memory traces on constant contralateral trials will also be able to decode contralateral-shifting Swap trials (orange). (C) The novel ensemble model posits that interhemispheric inputs activate a distinct ensemble (right) from visual inputs (left), even for the same memory trace. (D) It predicts failure of contralateral-trained decoders to generalize to contralateral-shifting trials. (E) For most of the post-saccadic delay, cross-decoding accuracy for ipsilateral-to-contralateral Swap trials (orange) did not significantly differ from constant ipsilateral trials, as predicted by the novel ensemble model. Near the end of the delay, a significant difference emerged (stars), indicating contralateral-shifting trials became more similar to constant contralateral trials, as predicted by the generic ensemble model. See also Figure S3.

We used a cross-classification method to adjudicate between the models. We trained a classifier to decode the object identity and its upper/lower location from population spiking at each time point on contralateral No-swap trials. We tested whether these classifiers could predict the same information on ipsilateral-to-contralateral Swap trials, which brought that same object to the same location as the contralateral No-swap trials. Note that training and testing were both performed on the same cortical hemisphere—separately for each hemisphere, then results were pooled across them—meaning the cross-classification is across *task conditions*, not cortical hemispheres. Thus, we tested whether the same information was reflected in the same neural pattern in a given hemisphere, regardless of how it arrived there. If both conditions activate the same ensembles, as assumed by the generic trace model (Fig. 6A), then this cross-classification (Fig. 6B, orange) should result in high decoding accuracy, similar to that obtained from both training and testing on constant contralateral trials (Fig. 6B, desaturated green). If these conditions activate different neural ensembles, as suggested by the novel ensemble model (Fig. 6C), then cross-classification accuracy (Fig. 6D, orange) should be poor, similar to that obtained from cross-classification testing on constant ipsilateral trials (Fig. 6D, desaturated brown). We find evidence for both models at different time points during the post-saccade delay.

For reference, we replot (from Fig. 4B) classification accuracy when both training and cross-validated testing were performed on constant contralateral trials (Fig. 6E, desaturated green). As a control, we computed cross-classification accuracy when the same No-swap contralateral trained classifiers were tested on No-swap ipsilateral trials (Fig. 6E, desaturated brown). This reflects baseline cross-classification generalization due solely to any bilaterality in the prefrontal neural code. This control analysis resulted in poor accuracy, further evidence for independence between the contralateral and ipsilateral hemifields. For most of the post-saccade delay, cross-classification of the contralateral-shift trials (Fig. 6E, orange) was also poor and not significantly different from the control (*p* ≥ 0.01, paired *t*-test). Here, classifier training and testing were performed at the same time points relative to the end of the delay. Similar results were obtained with training and testing at all possible relative times (Fig. S3), indicating results are not dependent on the specific timing scheme used.

These results mostly support the novel ensemble model. For much of the post-saccade delay, decoders generalized poorly from the constant contralateral trials to those in which it was transferred from the opposite hemisphere (Fig 6E). The initial bump in decoding accuracy seen when the saccade shifted the remembered location from ipsilateral to contralateral (Fig. 4B) was not detected using the classifiers from No-swap trials. Thus, different neural ensembles were activated by the same sensory information in WM, depending on whether it arrived via ipsilateral visual inputs or via the contralateral hemisphere. However, near the end of the delay, the cross-classification of contralateral-shift trials increased relative to the control (Fig. 6E, *p* < 0.01, paired *t*-test). This suggests that, in anticipation of using the WM, interhemispherically-transferred memory traces converged somewhat toward an ensemble representation similar to feedforward-induced traces.

### Interhemispheric synchrony during memory transfer

Our results thus far suggest transfer of WM traces between prefrontal hemispheres. If so, we should expect to see evidence of communication between hemispheres around the time of putative transfer. We would further expect signals to flow causally from the hemisphere contralateral to the initial sample hemifield (the “sender”) toward the hemisphere contralateral to the post-saccade hemifield (the “receiver”). Evidence suggests phase synchrony between cortical areas helps regulate the flow of information (Fries, 2015). Thus, we measured oscillatory synchrony between LFPs in the two prefrontal hemispheres using pairwise phase consistency (PPC), an unbiased measure of phase synchrony (Fig. 7).

**Figure 7.**
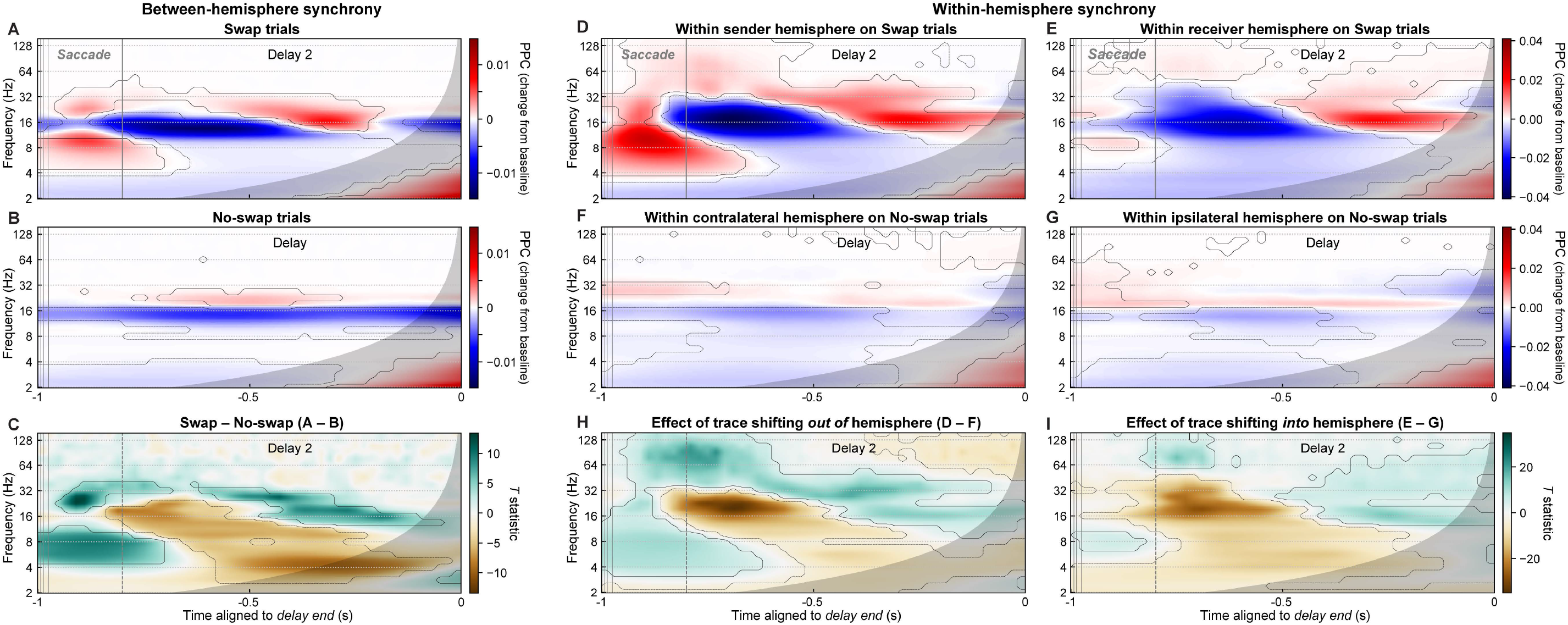
Interhemispheric beta/theta synchrony may mediate working memory trace transfer. (A,B) Mean phase synchrony (pairwise phase consistency, PPC) between all pairs of LFPs in the two prefrontal hemispheres, for Swap (A) and No-swap (B) trials, expressed as the change in PPC from the pre-sample fixation-period baseline. Contours indicate significant differences from baseline. Gray regions indicate time points with possible influence of test-period effects. (C) Contrast (paired t-statistic map) between Swap and No-swap conditions. Contours indicate significant between-condition difference. During the time period of putative interhemispheric memory trace transfer (–1 to –0.8 s), there was a significant increase (green) in interhemispheric synchrony in the theta (∼4–10 Hz) and beta (∼18–40 Hz) bands, and a decrease (brown) in the alpha/low-beta band (∼11–17 Hz). (D–E) PPC between LFP pairs within the sender (D) and receiver (E) hemisphere on Swap trials. (F–G) PPC between LFP pairs within the contralateral (F) and ipsilateral (G) hemispheres on No-swap trials. (H–I) T-statistic maps for contrasts between Swap and No-swap results for each hemisphere. These contrasts represent the effect of a WM trace shifting out of (H) and into (I) a hemisphere.

Around the saccade on Swap trials—when WM trace transfer putatively occurs— interhemispheric theta (∼4–10 Hz) and high-beta (∼18–40 Hz) synchrony both exhibited a transient peak (Fig. 7A; *p* < 0.01, paired *t*-test vs. pre-sample baseline). No such peaks were observed at analogous time points on No-swap trials (Fig. 7B). These differences were confirmed by examining the contrast between Swap and No-swap trials (Fig. 7C; *p* < 0.01, paired *t*-test). In contrast, during this period there was a suppression of interhemispheric synchrony relative to baseline within the alpha/low-beta band (∼11–17 Hz), which extended into the post-saccade delay period (Fig. 7C). Similar results, but with higher-frequency positive effects extending into the gamma band (∼40–100 Hz), were observed for synchrony between LFPs at distinct sites within each hemisphere (Fig. 7D–I). Theta and gamma increases were stronger in the sender hemisphere (Fig. 7D,F,H). Alpha/beta suppression from baseline—like beta power—was more similar in strength between hemispheres. These results suggest evidence for interhemispheric signal communication underlying memory trace transfer, and that communication occurs via theta and high-beta—but not alpha/beta—synchrony.

To test whether signals flow from the sender to the receiver hemisphere, we measured spectral Granger causality between LFPs in the two prefrontal hemispheres. Granger causality is a measure of how much variance in one signal can be explained by the recent history of another (here, LFPs at distinct sites), beyond what can be explained by the signal’s own dynamics. Spectral Granger expresses these causal interactions in the frequency domain (Dhamala et al., 2008). As predicted, causality was greater in the sender-to-receiver direction (Fig. 8A, green) than in the opposite, receiver-to-sender direction (orange). Though this difference was significant for all frequencies from approximately 10–40 Hz (Fig. 8B, gray stars; *p* < 0.01, paired *t*-test), it was greatest at the same high-beta frequencies that synchrony peaked at (∼20–40 Hz; Fig. 8B, gray curve). This asymmetric directionality was not observed in No-swap trials between sites contralateral and ipsilateral to the sample location (Fig. 8C–D). Between-region differences in signal-to-noise ratio (SNR) can result in false-positive Granger causality. Since our results were pooled across left-to-right and right-to-left hemispheres, any between-hemisphere SNR differences should have averaged out. To further rule out this possibility, we recomputed Granger causality on time-reversed data, which preserves SNR differences, but should reverse the direction of true causal signal flow (Bastos and Schoffelen, 2016; Haufe et al., 2013). This is exactly what we observed (Fig. S4). These results indicate that, during the time period of putative WM trace transfer, signals flow in the predicted direction, from the sender to the receiver hemisphere.

**Figure 8.**
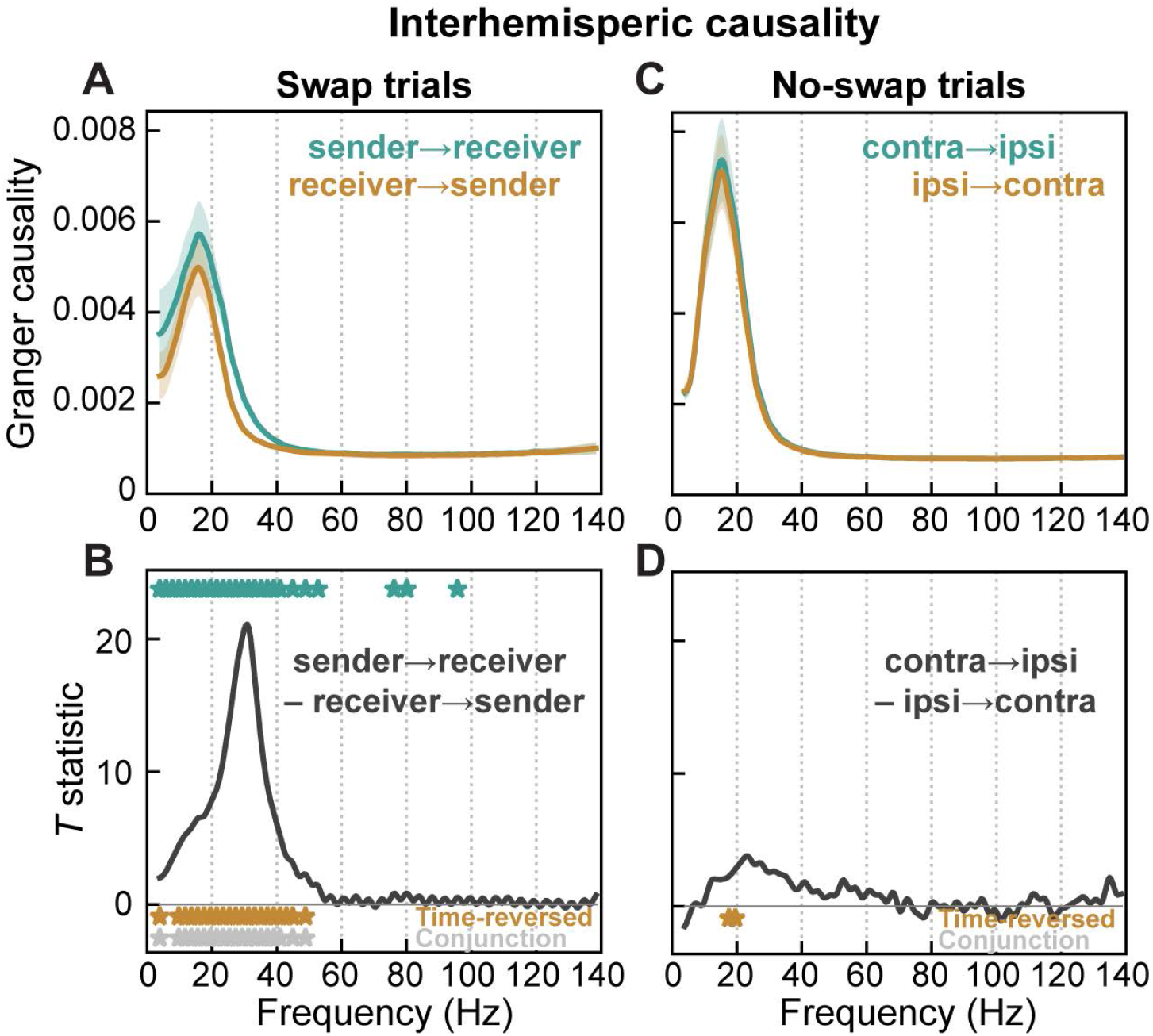
Granger causality indicates flow between prefrontal hemispheres in same direction as putative memory trace transfer. (A–B) Spectral Granger causality in Swap trials during time period of putative memory trace transfer from prefrontal hemisphere contralateral to initial sample location (“sender”) to hemisphere contralateral to post-saccade location (“receiver”). (A) Mean (± SEM) causality in the sender-to-receiver direction (green), and in the receiver-to-sender direction (orange). (B) T-statistic for contrast between causal directions. Causality was significantly greater in the sender-to-receiver direction across all frequencies ∼10–40 Hz (green stars). This directional asymmetry reversed, as expected, for time-reversed data (orange stars; full results in Figure S4). Gray stars indicate frequencies significant for both forward and time-reversed data. (C–D) No asymmetry in interhemispheric causality was observed during analogous time points (relative to delay end) in No-swap trials between contralateral and ipsilateral hemispheres.

## DISCUSSION

Our results suggest that WM traces can be transferred from one prefrontal hemisphere to the other. Previous studies have only provided indirect evidence of this. In WM tasks, when sample and test stimuli appear in opposite visual hemifields, masking during the delay is more effective in the expected test hemifield (Zaksas et al., 2001). Associative cues presented to one visual hemifield can elicit associated representations in the opposite-hemisphere visual cortex (Tomita et al., 1999). Here, we provided direct neurophysiological evidence of WM transfer between hemispheres.

A possible anatomical substrate is direct connections between hemispheres via the corpus callosum (Barbas and Pandya, 1984). Cortical feedforward processing has been shown to be mediated by gamma and theta oscillations while feedback processing is mediated by alpha/beta oscillations (Bastos et al., 2015; Buschman and Miller, 2007; van Kerkoerle et al., 2014). Our results showed increases in interhemispheric theta and high-beta synchrony, and decreases in alpha/low-beta, during the presumed time of memory trace transfer. One possibility is that interhemispheric communication occurs both in frequency bands utilized by feedforward (theta) and feedback (beta) processing. Perhaps this allows flexible interaction of interhemispheric signals with either feedforward or feedback signals. Another possibility is that higher frequency cortical communication—typically in the traditional gamma band—is shifted slightly downward in frequency (to 40 Hz and lower) to maintain reliable transmission despite the time delays imposed by long callosal axons. Of course, there are other possible routes for interhemispheric transfer other than direct prefrontal callosal connections. Signals might be transferred at some higher-level region then conveyed via feedback to PFC, or conceivably transfer could happen at lower levels and feed forward to PFC. Ultimately, resolving this issue will likely require causal manipulation of connectivity.

Gamma and alpha/beta oscillations have characteristic WM-related temporal dynamics— gamma is elevated and beta suppressed during encoding (sample period) and read-out (end-of-delay “ramp-up”) of WM, while the opposite is true during WM maintenance (delay period; Lundqvist et al., 2016, 2018). We replicated these anti-correlated gamma/beta dynamics both when a WM is statically maintained, and when it is transferred between hemispheres. We found that theta dynamics also broadly correlate with gamma and anti-correlate with alpha/beta. We also found similar dynamics in these frequency bands in the synchrony between distinct sites both within a prefrontal hemisphere and between hemispheres, with the exception that synchronization between hemispheres at traditional gamma frequencies was absent.

To better understand the temporal dynamics of interhemispheric WM transfer, we compared the latencies of all studied neural signals (Fig. S5). The earliest signals after the saccade instruction included beta synchrony within the sender hemisphere and between hemispheres, consistent with the idea that it may be involved in establishing a communication channel for interhemispheric transfer. Beta signals then exhibited a bilateral decrease around the time the receiver hemisphere begins to convey information about the WM trace, consistent with the idea that suppressing beta may disinhibit cortex and allow WMs to be expressed (Lundqvist et al., 2016). Finally, WM information decreased in the sender hemisphere about 120 ms after the rise in the receiver. Interhemispheric transfer in this task thus involves a “soft handoff”, in which information overlaps in time in both hemispheres before being cleared out of the sending hemisphere. This is similar to what has been found for tracking moving objects across the midline (Drew et al., 2014).

Another way to think about our results is in terms of reference frames. In a retinotopic (gaze-centered) reference frame, locations are described relative to gaze (Fig. S6A, top). Virtually all studied areas of visual cortex code in a retinotopic reference frame (Fig. S6B, top; Cohen and Andersen, 2002; Golomb and Kanwisher, 2012). In contrast, a spatiotopic (world-centered) reference frame represents locations in the real-world, independent of gaze (Fig. S6A,B, bottom). In addition to other more established non-retinotopic spatial coding schemes (Chafee et al., 2007; Graziano and Gross, 1998; Olson, 2003), explicit spatiotopic reference frames may exist in higher-level cortex (Dean and Platt, 2006; Duhamel et al., 1997).

In our study, whether the eyes were fixated to the left or right, the sample item was always at the same central spatiotopic (real-world) location. Thus, if prefrontal cortex encoded locations in a spatiotopic reference frame (Fig. S6B, bottom), its activity should be relatively invariant to the location of the sample object relative to the eyes (Fig. S6C, bottom row). Arguing against this possibility is the differentiation between contralateral and ipsilateral sample locations in PFC activity (Fig. 2). This suggests a retinotopic reference frame (Fig. S6C, top row). On the other hand, if WM—at the cognitive level—also maintained locations in a retinotopic reference frame, remembered locations would be anchored to the location of gaze and would simply shift around with a saccade (Fig. 9A, top). This makes predictions (Fig. 9C, left column) identical to the stable trace model (Fig. 2A,C), i.e., that the memory trace remains in the original hemisphere.

Instead, we found that neural signatures of laterality invert after the saccade (Fig. 4–5), ruling out a retinotopic reference frame for WM (Fig. 9C, left column). Our results are consistent with the remembered location being maintained within a spatiotopic reference frame at the cognitive level (Ong et al., 2009), but within a retinotopic reference frame at the neural level (Fig. S6C, upper-right quadrant).

How can we reconcile this? One possibility is that cognition reflects some putative higher-level area with a spatiotopic reference frame that updates the retinotopic representation after each saccade. Arguing against this is the relative paucity of evidence for any explicit spatiotopic representation in the visual cortical hierarchy (Dean and Platt, 2006; Duhamel et al., 1997; Golomb and Kanwisher, 2012). Alternatively, spatiotopic cognition may not rely on an explicit spatiotopic neural representation. It could instead be coded implicitly by a retinotopic representation that is locally updated after a saccade. This would shift a remembered location in the opposite direction of each saccade vector to its new coordinates on the retinotopic map (Pouget and Snyder, 2000). For a midline-crossing saccade, this would entail transfer of the memory trace between the left and right visual hemifields/hemispheres.

In many areas of visual and visuomotor cortex, receptive fields (RFs) show anticipatory shifts to their future, post-saccadic location even before the onset of a saccade (Colby et al., 1995). Other studies suggest that during a saccade, RFs contract toward the location of the saccade target then later expand out to their final post-saccadic location (Chen et al., 2018; Neupane et al., 2016; Zirnsak et al., 2014). This implies a change in the population neural code for location around the time of a saccade because the ensemble responsive to a given location during the RF contraction will be different from those responsive to the same location before and after the saccade. This could explain the lack of cross-decoding between static and shifted memory traces (Fig. 6C). These dynamics have yet to be demonstrated in PFC but their properties in other areas argue against a role. RF contraction effects extend to only ∼300 ms after a saccade in area V4 (Neupane et al., 2016) and the frontal eye field (Chen et al., 2018), whereas poor cross-decoding extends to over 500 ms in our results. Contraction effects in V4 occur mainly for neurons with RF in the *same* hemifield as the saccade endpoint (Neupane et al., 2016), whereas in the key ipsilateral-to-contralateral condition in our results, RFs would be in the *opposite* hemifield to the saccade.

An alternative account is motivated by findings of “nonlinear mixed selectivity” in PFC (Rigotti et al., 2013). It suggests PFC is best understood as a random network, in which unique ensembles are activated by different combinations of input features and task contexts (Bouchacourt and Buschman, 2019) . This predicts distinct ensembles are activated depending on the route by which information arrives, feedforward vs. interhemispheric. Regardless of the specific mechanism, our results indicate that just before the memory trace is to be read out for comparison with a test object, its neural code shifts to become more like the ensemble used for static, feedforward-induced memory traces in the same location. This convergence likely facilitates downstream comparison and decision-making processes by allowing similar mechanisms and read-out weights for reading the same information out from WM stores.

Processing in most visual cortical areas, including PFC, is strongly biased toward the contralateral visual hemifield (Funahashi et al., 1990; Hagler and Sereno, 2006; Kastner et al., 2007; Medendorp et al., 2007; Pasternak et al., 2015; Rainer et al., 1998; Voytek and Knight, 2010; Wimmer et al., 2016). Prefrontal spiking activity (Buschman et al., 2011) and gamma power (Kornblith et al., 2015) increase with WM load—the number of items held in memory at one time—but only for items in the contralateral visual hemifield. In contrast, beta power shows increasing suppression for increasing numbers of items in either visual hemifield (Kornblith et al., 2015; Medendorp et al., 2007). Our results confirm these findings. This distinction might reflect the fact that beta oscillations are thought to correlate with broadly-selective inhibitory processes (Engel and Fries, 2010; Jensen and Mazaheri, 2010; Lundqvist et al., 2016). It might also reflect beta having a stronger influence from more bilateral top-down or recurrent signals, while spiking and theta/gamma oscillations are dominated by feedforward signals from strongly lateralized visual cortex (Bastos et al., 2015; van Kerkoerle et al., 2014).

A somewhat surprising result of this lateralization is that WM capacity is largely independent between the two visual hemifields. WM has a very limited capacity for holding multiple items at one time (Luck and Vogel, 1997, 2013). However, in both monkeys (Buschman et al., 2011) and humans (Delvenne, 2005; Umemoto et al., 2010), even when capacity is saturated in one visual hemifield, additional items can be stored in WM if they appear in the opposite hemifield. Similar effects of hemifield independence have been observed with spatial attention (Alvarez et al., 2012) and attentional tracking of moving objects (Alvarez and Cavanagh, 2005). This strong hemifield independence, however, seems inconsistent with the apparently seamless nature of visual WM. Our results provide a possible resolution to this paradox. They suggest that, in such situations, the two prefrontal hemispheres briefly sync up using theta and beta oscillations in order to physically transfer a WM trace from one cortical hemisphere to its new location on the retinotopic map in the opposite hemisphere.

We found neural dynamics that could support the seamless transfer of information between the cerebral hemispheres. We expect some of these same mechanisms are also employed when interhemispheric communication is required for other cognitive processes, such as tracking moving objects across the midline or comparing visual information between the left and right hemifields. Fast, reliable interhemispheric communication is critical for many real-world behaviors, including sports, driving, and air traffic control. Interhemispheric communication is also thought to be disrupted in some disorders, such as dyslexia (Dhar et al., 2010). We hope that an understanding of the neural mechanisms of interhemispheric communication may lead to new ways to repair and optimize it.

## ACKNOWLEDGEMENTS

We thank Alexa D’Ambra for assistance, and Jesus Ballesteros, Andre Bastos, Sayak Bhattacharya, Alex Major, Morteza Moazami, Dimitris Pinotsis, and Jefferson Roy for helpful comments. This work was supported by NIMH R37MH087027, ONR MURI N00014-16-1-2832, The JPB Foundation (E.K.M.), and NIGMS T32GM007753 (J.A.D).

## AUTHOR CONTRIBUTIONS

J.A.D., S.K., and E.K.M. designed the experiments. J.A.D., M.K.M., and S.K. performed the experiments and recorded the data. S.L.B. curated the data and conceived and performed the analyses. S.L.B, E.K.M., and M.L. wrote the manuscript.

## DECLARATION OF INTERESTS

The authors declare no competing interests.

## STAR METHODS

### RESOURCE AVAILABILITY

#### Lead Contact

Requests for information should be directed to the lead contact, Earl Miller.

#### Materials Availability

This study did not generate new unique reagents.

#### Data and Code Availability

The neural data and code used in this study will be made available upon reasonable request to the lead contact.

## EXPERIMENTAL MODEL AND SUBJECT DETAILS

The nonhuman primate subjects in our experiments were two adult (ages 16 and 9) rhesus macaques (*Macaca mulatta*), one male and one female. All procedures followed the guidelines of the Massachusetts Institute of Technology Committee on Animal Care and the National Institutes of Health.

## METHOD DETAILS

### Behavioral paradigm

Subjects performed a delayed nonmatch-to-sample working memory (WM) task (Fig. 1A). They began task trials by holding gaze for 700 ms on a fixation point randomly displayed at 4.5° left or right of the center of a computer screen. A sample object was then shown for 700 ms in the center of the screen, thus in the right or left visual hemifield, respectively. Two sample objects were used each session, chosen from a commercial photo library (Hemera Photo-Objects). Sample objects were displayed in one of two positions, 3.4° above or below the screen center. After a 1.6 s delay, a test object was displayed for 400 ms. The monkeys were required to saccade to it if it did not match the remembered sample in either object identity or upper/lower location. If the test was identical to the sample, they withheld response. Then, after a 100 ms blank period, a nonmatching test object was always shown, which required a saccade. Response to the non-match was rewarded with juice, followed by a 3.2 s inter-trial interval.

A random 50% of trials had an uninterrupted 1.6 s WM delay (Fig. 1A, left). In the other 50%, at 800 ms into the delay, the fixation point jumped to the opposite location on the screen (Fig. 1A, right). The monkeys were trained to immediately saccade to it and reacquire fixation. Once fixation was acquired again, the WM delay was continued for another 800 ms, equating the full time of fixated delay period with the No-swap condition.

All stimuli were displayed on an LCD monitor. An infrared-based eye-tracking system (Eyelink 1000 Plus, SR-Research, Ontario, CA) continuously monitored eye position at 1 kHz.

### Electrophysiological data collection

The subjects were chronically implanted bilaterally in the lateral prefrontal cortex (PFC) with four 8×8 iridium-oxide “Utah” microelectrode arrays (1.0 mm length, 400 µm spacing; Blackrock Microsystems, Salt Lake City, UT), for a total of 256 electrodes (Fig 1C). Arrays were implanted bilaterally, one array in each ventrolateral and dorsolateral PFC. Electrodes in each hemisphere were grounded and referenced to a separate subdural reference wire. LFPs were amplified, low-pass filtered (250 Hz), and recorded at 30 kHz. Spiking activity was amplified, filtered (250– 5,000 Hz), and manually thresholded to extract spike waveforms. All threshold-crossing spikes on each electrode were pooled together and analyzed as multi-unit activity (MUA).

## QUANTIFICATION AND STATISTICAL ANALYSIS

All correctly performed trials were included in analyses. All analyses of individual MUA and LFP channels were averaged across all electrodes in each hemisphere, and analyses of channel pairs were averaged across all between-hemisphere or all within-hemisphere pairs. Analysis was initially performed separately for each prefrontal hemisphere and sample object hemifield. Results were then pooled based on whether the sample was contralateral or ipsilateral to a given hemisphere by averaging across appropriate hemisphere/hemifield combinations. This resulted in a set of observations for each experimental session. All plots depict means and standard errors across all 56 sessions in this dataset, and all statistics were performed with sessions treated as observations (*n* = 56). All preprocessing and analysis was performed in Python 3.6 or Matlab R2019b (The Mathworks, Inc, Natick, MA).

### Preprocessing

Spike rates were computed by binning spike timestamps in non-overlapping 50 ms windows. Spike rates were square-root transformed prior to analysis to convert their Poisson-like distributions to approximately normal. LFPs were re-referenced offline to remove any common-source noise by subtracting off the mean across all electrodes in each array. Evoked potentials were removed by subtracting off the mean signal across trials within each condition (object, upper/lower location, and visual hemifield). Thus, all of our analysis is on the remaining induced component. For most analyses, the resulting signals were convolved with a set of complex Morlet wavelets (wavenumber 6). LFP power was log-transformed to render its distribution approximately normal.

### Mean activity analysis

To normalize out any overall differences in activity between neurons and task conditions, we z-scored spike rates to the fixation baseline. Rates were mean-pooled across the 200 ms before sample object onset separately for each neuron and condition, and the mean and standard deviation across all within-condition trials was computed. These were used to z-score rates across all time points and trials for that neuron and condition. The same transformation was used for analysis of LFP power, except it was also computed separately for each frequency, in order to also normalize out the typical 1/*f* distribution of power across frequency.

To assay for spiking activity changes from baseline in either direction (increases or decreases), we took the absolute value of the measured z-scores for each multi-unit (Fig. S1A). Significance was evaluated using paired-observation tests relative to baseline. We also measured delay activity separately for subpopulations of multi-units with responses above or below baseline (Fig. S1B). Since PFC units often have complex dynamics, this segregation into subpopulations was performed independently for each unit and time point. To avoid circularity in the analysis, segregation into subpopulations and computation of mean activity within each subpopulation were performed on odd and even-numbered trials, respectively.

### Population decoding analysis

Spike rates for all multi-units within a given hemisphere were used as independent features in a linear classifier that decoded which of four task conditions—two objects × two upper/lower locations—was present in each trial. Classification was performed independently on spike rate data from each time point (50 ms window). All reported classification accuracies were obtained via 5-fold cross-validation, in which trials were randomly split into five non-overlapping subsets and each classifier was trained on four of these, while its accuracy was evaluated on the final, untrained one. This process was repeated five times with each subset acting as the test set once, and the final results were averaged across the five folds. The same procedure was used for the cross-classification analysis (Fig. 6), except that training and testing trials were selected from different task conditions. For the cross-temporal analysis (Fig. S3), cross-classification was also performed using all possible combinations of pairs of time points for training and testing. All decoding analysis was performed with a linear discriminant classifier with optimal covariance shrinkage (Ledoit and Wolf, 2003), using the Python scikit-learn library.

As an alternative measure of difference in neural activity elicited by task conditions, we also computed the mean rate for each multi-unit’s preferred, intermediate, and nonpreferred WM items (objects × upper/lower locations; Fig. S1C–D). This was computed separately for each multi-unit and time point, since PFC units often have complex dynamics with different preferences at different time points. To avoid circularity, calculation of item preference and of mean rates for the preference-sorted items was performed on odd and even-numbered trials, respectively.

### Synchrony analysis

LFP-LFP phase synchrony was computed from the phase of the complex wavelet transform, using the pairwise phase consistency (PPC). PPC is a measure of how consistent across trials the relative phase angles between two signals are, independent of their absolute phase. It is an unbiased estimator of the square of the mean vector resultant length (Kornblith et al., 2015; Vinck et al., 2010). Presented results represent averages across all electrode pairs between hemispheres (Fig. 7A–C), all electrode pairs within the hemisphere contralateral to the sample hemifield (the “sender” on Swap trials; Fig. 7D,F,H), and all electrode pairs within the hemisphere ipsilateral to the sample hemifield (“receiver” on Swap trials: Fig. 7E,G,I) within each session. These values were then averaged across all sessions.

### Causality analysis

Directional causal influences between LFPs in the two prefrontal hemispheres were measured using bivariate nonparametric spectral Granger causality (Dhamala et al., 2008). Traditional Granger causality quantifies how much variance in one signal can be explained by the recent history of another signal, beyond what can be explained by the history of the signal itself. Spectral Granger expresses these causal interactions in the frequency domain. Unlike traditional parametric causality measures, it is estimated directly via factorization of the cross-spectral density matrix, without relying on estimation of a specific autoregressive model. For this analysis only, LFP spectra were computed around the putative trace transfer period (–1 to –0.5 s relative to delay end) via the multitaper method (4 Hz frequency bandwidth, 3 dpss tapers). Trials were balanced across Swap and No-swap conditions in each session by sampling a random subset of trials from the larger group. Presented results represent averages across all between-hemisphere electrode pairs within each session, and across all sessions. As a control, this analysis was also performed on data reversed in time (Fig. S4). This is expected to reverse the direction of true causality, but preserve the direction of any apparent causality due to confounding between-area differences in power or signal-to-noise (Bastos and Schoffelen, 2016; Haufe et al., 2013). Inference was based on the conjunction of significance in the standard analysis and significance in the opposite causal direction in the time-reversed analysis. This analysis was performed using the FieldTrip toolbox for Matlab (Oostenveld et al., 2011).

### Oscillatory burst analysis

Methods followed those of our previous work on oscillatory bursts (Lundqvist et al., 2016). Power was computed from a wavelet transform, as described above, and pooled within several frequency “sub-bands”: low-beta (10–20 Hz), high-beta (20–32 Hz), low-gamma (40–65 Hz), gamma (55–90 Hz), and high-gamma (70–100 Hz). Within each sub-band, the mean and SD of power across all trials and within-trial time points was computed and used to threshold the data at 2 SDs above the mean. Bursts were defined as periods where power remained above threshold for at least 3 oscillatory cycles (defined at the center frequency of each sub-band). This resulted in a binary representation of time points within vs. outside bursts. These were pooled separately within all beta and gamma sub-bands by labelling as bursts time points where a burst occurred within any sub-band (logical OR across sub-bands). Burst rate for each task condition, band, and time point was computed as the proportion of trials containing within-burst time points. Finally, the difference in burst rate at each time point from the fixation baseline was computed and plotted in Fig. S2.

### Latency analysis

Latencies were measured for all reliable temporal features—points halfway from a baseline to a peak or trough—of neural signals surrounding the epoch of WM trace transfer (Fig. S5). These were measured on across-session mean neural signals, and their 99% confidence intervals were computed by bootstrapping across sessions 10,000 times. Each bootstrap iteration resampled sessions with replacement, computed the same across-session mean neural signals, and estimated resampled latencies from them.

### Postprocessing and plotting

To clarify trends in the data, all plotted results were smoothed with 1D or 2D Gaussian kernels for plotting purposes only. The Gaussian standard deviations used were 10 ms for all time axes, and 0.05 octaves for all frequency axes.

The temporal extent over which time-frequency effects due to the test object could possibly extend into the delay period was computed as the time for wavelet power at each frequency to drop by a factor of *e*^−2^ relative to the test stimulus onset (Torrence and Compo, 1998). This is a conservative overestimate of the backward extent of test-object effects since it does take any latency of test-evoked neural signals into account. This period is plotted as a transparent gray region at the right of all time-frequency and frequency-band summary plots.

Due to off-axis diagonal structure in some of the time-frequency data (time-frequency inseparability), it cannot be fully captured in any 1D summary. Thus, all statistical analyses and inference were performed on the full time-frequency data. However, we also include 1D plots of the time course within frequency bands simply to aid in understanding the results. The cutoffs for each frequency band were chosen to approximate values typically used to delineate standard frequency bands in the literature: theta (3–8 Hz), beta (10–32 Hz), and gamma (40– 100 Hz).

### Hypothesis testing

All hypothesis tests used non-parametric randomization methods that do not rely on specific assumptions about data distributions (Manly, 2007). Each session was treated as an observation, and randomizations were performed across sessions. All randomization statistics were resampled 10,000 times and evaluated in a two-tailed fashion.

To test whether a mean value differed significantly from baseline, we used a randomized sign test in which a *t*-statistic was computed on both the observed data, and on data where the sign of each baseline-centered observation was randomly flipped. To test whether the means of paired observations were significantly different, we used a permutation paired *t*-test, in which a paired-sample *t*-statistic was computed on the observed data, and on data with the labels of each pair of observations randomly swapped. To test significance of multiple main effects and their interaction, we used a permutation 2-way ANOVA in which an *F*-statistic was computed on the observed data, and on data where the multi-factor labels were randomly shuffled as a group across trials. All tests were corrected for multiple comparisons across time points and/or frequencies using a procedure that controls the false discovery rate under arbitrary dependence assumptions (Benjamini and Yekutieli, 2001) using the Python statsmodels module.

## Notes

### Competing Interest Statement

The authors have declared no competing interest.

### Summary of Updates

Updated analyses and figures. Paper now “in press” at Neuron

