## Supplemental figures for "Interhemispheric transfer of working memories"

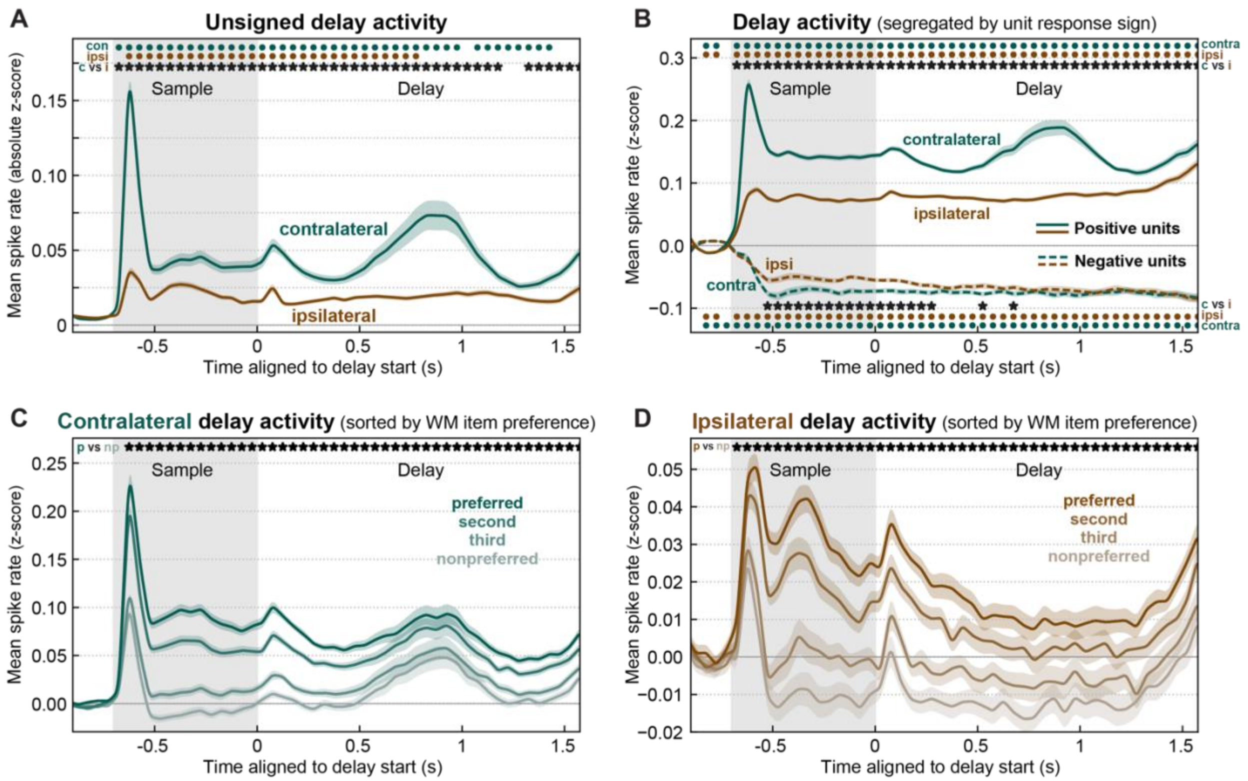


***Figure S1. Ipsilateral delay activity is balanced between enhancement and suppression; contralateral delay activity is biased toward enhancement (related to Figure 2A,B).*** *(A) Mean (± SEM) unsigned delay activity (absolute z-score of spike rates relative to baseline) for sample objects in the contralateral (green) and ipsilateral (brown) hemifields. In contrast to signed delay activity (Fig. 2A in main text), ipsilateral samples elicit significant absolute changes from baseline during the delay period (brown dots), though they remained significantly weaker (stars) than those elicited by contralateral samples. (B) Mean (± SEM) signed delay activity computed separately for units with cross-validated positive (solid lines) and negative (dashed lines) deviations from baseline. While contralateral samples (green) elicit stronger deviations in positive than negative units, ipsilateral samples (brown) elicit approximately balanced activity in both subpopulations, consistent with the near-baseline overall activity observed in main text Fig. 2A. (C–D) Mean (± SEM) signed delay activity for contralateral (C) and ipsilateral (D) samples, computed separately for each unit's cross-validated preferred (most saturated colors), intermediate, and nonpreferred (most desaturated colors) sample items (object and upper/lower location). Both hemifields elicit clear differences between preferred and nonpreferred items (stars), consistent with the decoding accuracy results in the main text (Fig. 2B). For ipsilateral samples (D), however, preferred items elicited above-baseline activity and nonpreferred items suppressed activity below baseline, resulting in near-baseline delay activity when pooled together (main text Fig. 2A).*

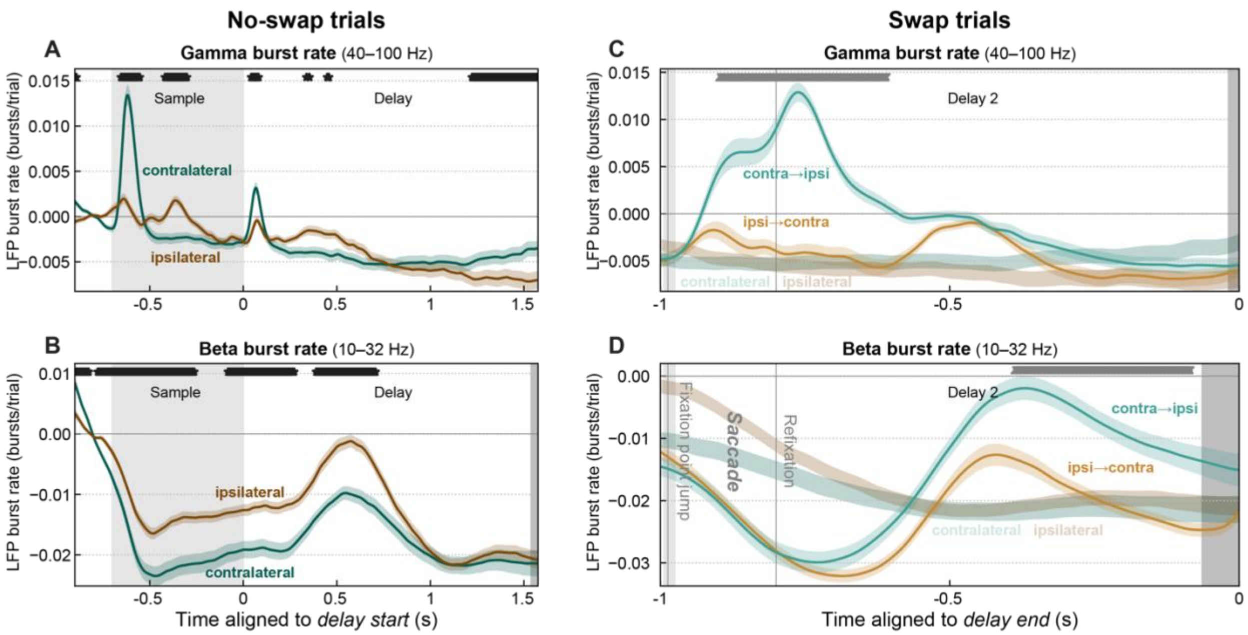


***Figure S2. Oscillatory burst rates produce results similar to oscillatory power (related to Figures 2F–H and 5D–F).*** *(A–B) Mean (± SEM) rate of oscillatory LFP gamma (A) and beta (B) bursts in No-swap trials for contralateral (dark green) and ipsilateral (brown) sample objects, expressed as difference from pre-sample baseline rate. Black stars: significant difference between contralateral and ipsilateral. Contralateral bias, temporal dynamics, and antagonistic gamma/beta relationship are broadly similar to that observed in LFP power (main text Fig. 2). One notable difference is that the contralateral bias for beta burst rates (greater suppression for contralateral than ipsilateral samples in B) is stronger and more extended into the delay than for beta power. (C–D) Gamma (C) and beta (D) burst rates on Swap trials. Gray symbols: significant sample hemifield × Swap/No-swap interaction. Results are broadly similar to those for oscillatory power (main text Fig. 5), though only beta burst rates show clear evidence of post-saccade inversion. Gray regions at right of all plots indicate time points that could be affected by temporal smearing of test-period effects.*

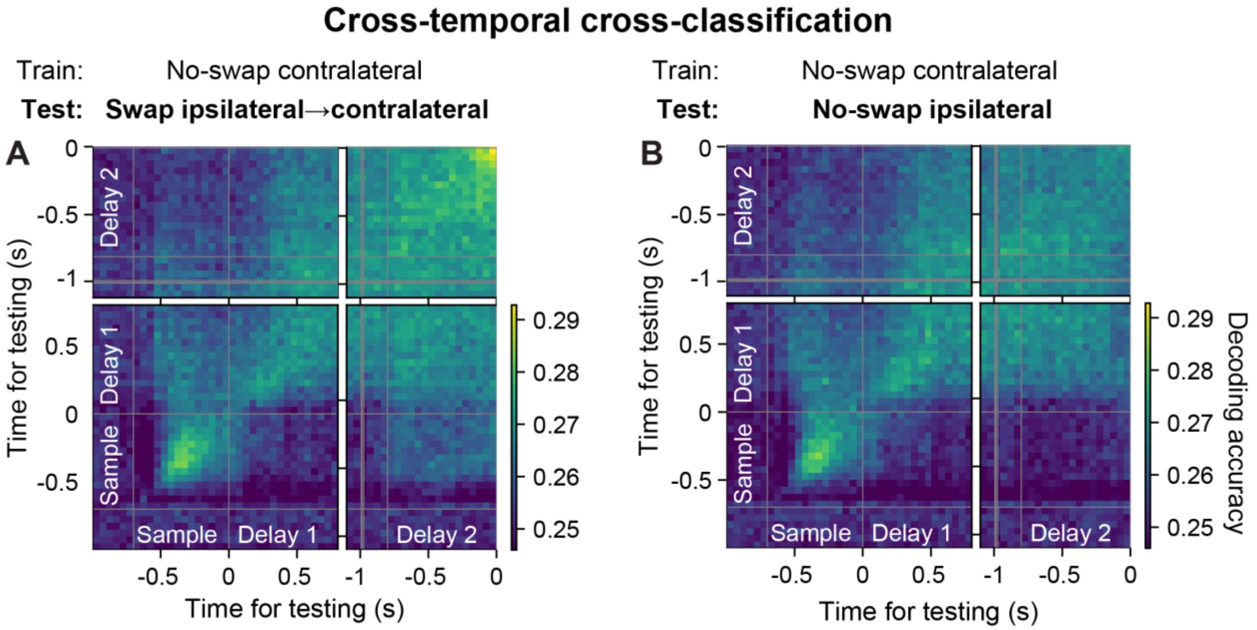


***Figure S3. Poor cross-decoding of contralateral-shifting trials at all relative times except for pre-test ramp-up (related to Figure 6).*** *(A) Mean decoding accuracy for classifiers trained on constant contralateral trials and tested on ipsilateral-to-contralateral Swap trials, for all combinations of training time (y-axis) and testing time (x-axis). (B) Cross-temporal decoding accuracy for classifiers trained on constant contralateral trials and tested on constant ipsilateral trials. Main diagonals correspond to training and testing at same time within trials (as plotted in Fig. 6E). This analysis confirms that cross-decoding of contralateral-shifting trials (A) is similar to the baseline provided by prediction of constant ipsilateral trials (B) at all training/testing times, except for the "ramp-up" at delay end (upper-right corner of A).*

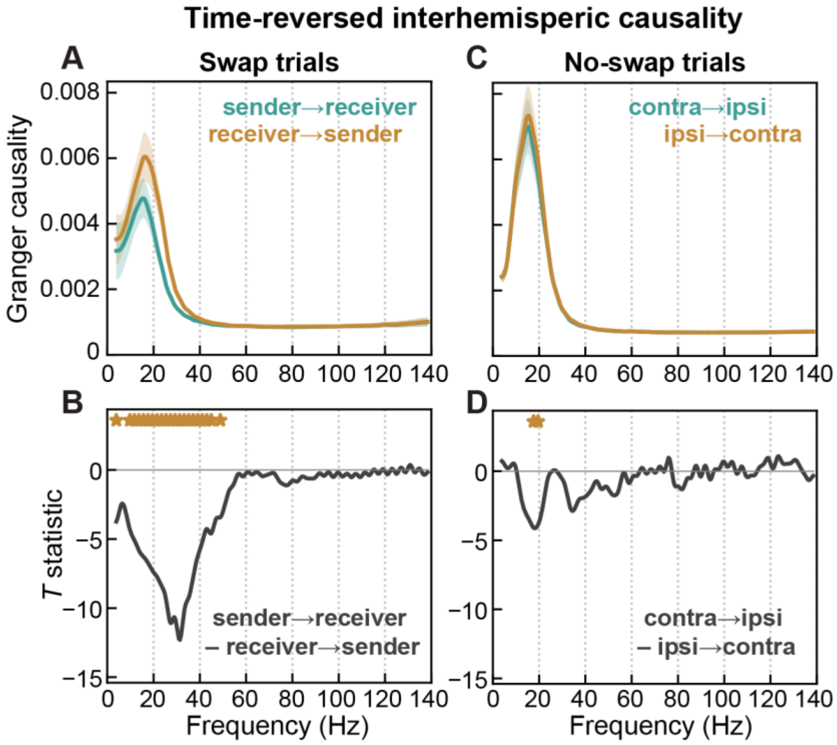


***Figure S4. Time-reversed analysis controls for signal-to-noise explanation of causality (related to Figure 8).*** *Spectral Granger causality on time-reversed data during Swap (A–B) and No-swap (C–D) trials. Time reversal preserves any differences in power or signal-to-noise ratio (SNR) between signals but reverses their temporal precedence. Thus, time reversal should also reverse the direction of true causal influences, while apparent causality due to power or SNR differences should be unchanged. Compared to causality measured on the original data (main text Fig. 8), time reversal indeed reversed the direction of causal flow—now the receiver-to-sender direction predominates (A,B). This control analysis indicates the observed Granger causality reflects true causal signal flow from the sender to receiver hemisphere, not differences in power or SNR between them.*

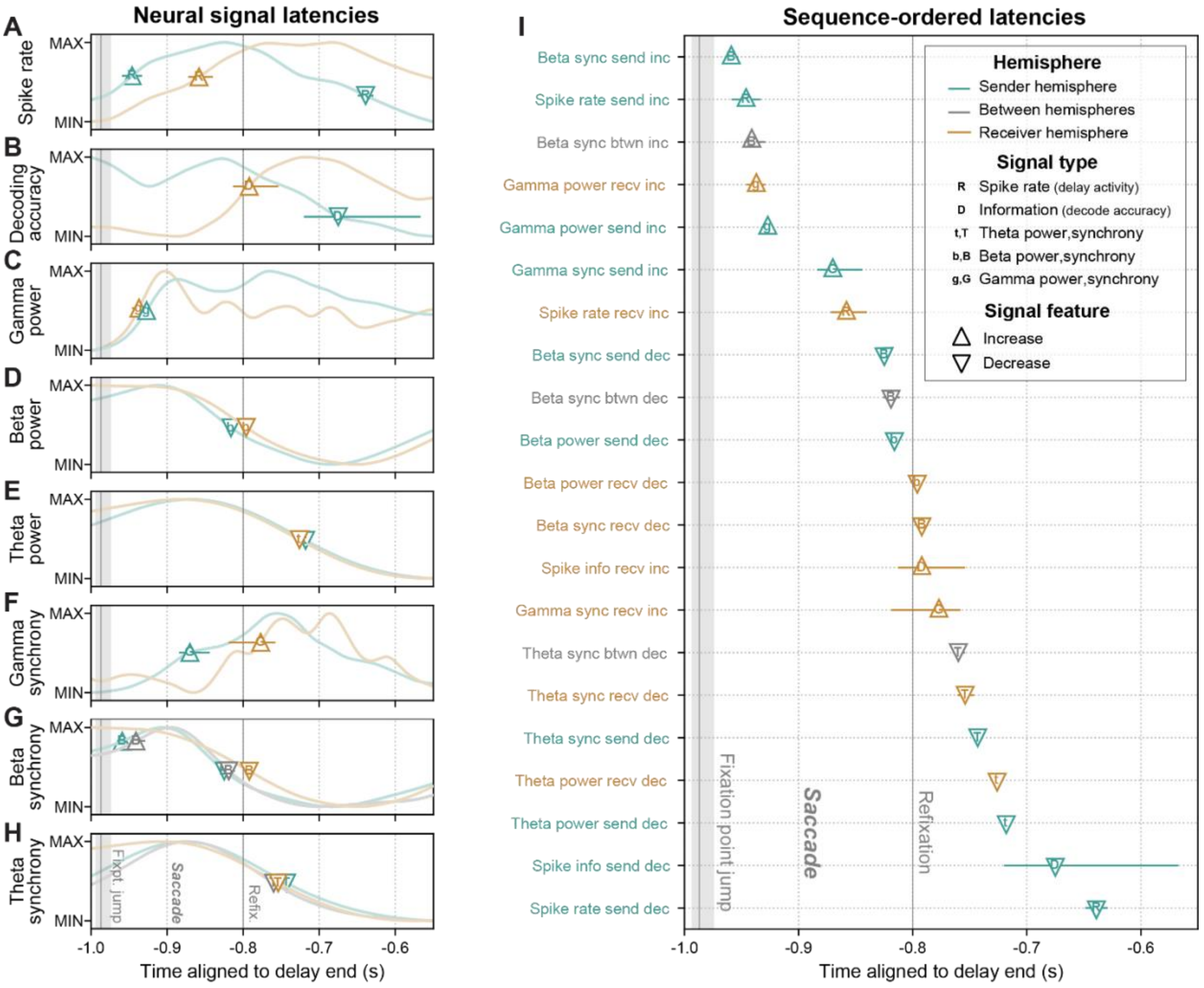


***Figure S5. Temporal sequence of neural signals during interhemispheric working memory trace transfer (related to Figures 2 and 7).*** *(A–H) Latencies (± 99% confidence interval) of temporal features (increases and decreases) of all studied neural signals, overlaid on the source signal: spike rate (A) and decoding accuracy (B); LFP gamma power (C), beta power (D), theta power (E), gamma synchrony (F), beta synchrony (G), and theta synchrony (H). Color indicates prefrontal hemisphere the signal was derived from, text symbols indicate signal type, and surrounding marker shapes indicate signal temporal feature (legend in I). Note time scale zoomed into period of putative WM trace transfer. (I) Latencies for neural signals of all hemispheres, signal types, and signal features reordered from earliest to latest. The earliest signals included increases in beta synchrony within the sender hemisphere and between hemispheres, consistent with a role for beta synchrony in establishing interhemispheric communication. At post-saccade refixation, beta signals decreased bilaterally as information about the WM increased in the receiver hemisphere. Finally, WM information decreased in the sender, ~120 ms after the increase in the receiver, evidence for a "soft handoff" of information between hemispheres.*

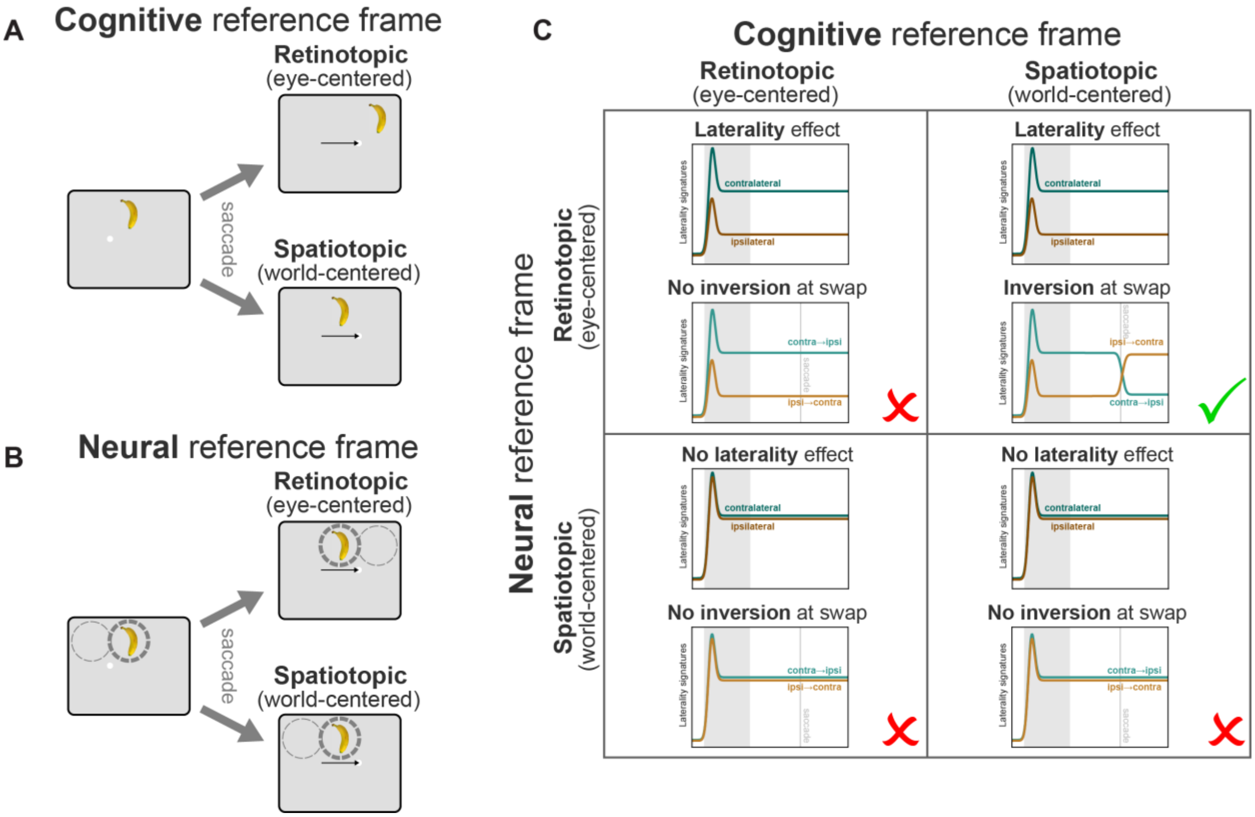


***Figure S6. Spatial reference frame interpretation of results (related to Figures 2, 4, 5, and Discussion).*** *(A) Illustration of two possible spatial reference frames for remembered locations in cognitive working memory. Left: An object in the right hemifield is encoded into working memory. Right: Effects of a saccade across the midline (black arrow) on the remembered location. Under a retinotopic reference frame (top), the remembered location shifts with the saccade. Under a spatiotopic reference frame (bottom), the remembered location remains anchored to its real-world location. (B) Illustration of two possible reference frames for neural representation of object location. Left: An object in the right hemifield activates the rightmost of two receptive fields (RFs; dashed circles). Right: Under a retinotopic reference frame (top), RFs shift with the eyes, and the object now activates the leftmost RF. Under a spatiotopic reference frame (bottom), RFs are anchored to locations in the real world, and the object continues to activate the rightmost RF. (C) Predictions of all combinations of cognitive (columns) and neural (rows) reference frames for our results. A spatiotopic neural reference frame (bottom row) predicts no change in activity across gaze positions, and thus no laterality effect in our data. This is inconsistent with our results (Fig. 2), ruling out these possibilities. A retinotopic cognitive reference frame (left column) predicts the remembered location shifts with the mid-delay saccade, and thus no inversion of neural laterality signatures. This is inconsistent with our results (Fig. 4–5), ruling out this possibility. Thus, a retinotopic neural reference frame, in conjunction with a spatiotopic cognitive reference frame (upper-right) is the only option consistent with our results.*
